# Development of a sequence-based reference physical map of pea (*Pisum sativum* L.)

**DOI:** 10.1101/518563

**Authors:** Krishna Kishore Gali, Bunyamin Tar’an, Mohammed-Amin Madoui, Edwin van der Vossen, Jan van Oeveren, Karine Labadie, Helene Berges, Abdelhafid Bendahmane, Reddy V.B. Lachagari, Judith Burstin, Tom Warkentin

## Abstract

Whole genome profiling (WGP) is a sequence-based physical mapping technology and uses sequence tags generated by next generation sequencing for construction of bacterial artificial chromosome (BAC) contigs of complex genomes. The physical map provides a framework for assembly of genome sequence and information for localization of genes that are difficult to find through positional cloning. To address the challenges of accurate assembly of the pea genome (~4.2 GB of which approximately 85% is repetitive sequences), we have adopted the WGP technology for assembly of a pea BAC library. Multi-dimensional pooling of 295,680 BAC clones and sequencing the ends of restriction fragments of pooled DNA generated 1,814 million high quality reads, of which 825 million were deconvolutable to 1.11 million unique WGP sequence tags. These WGP tags were used to assemble 220,013 BACs into contigs. Assembly of the BAC clones using the modified Fingerprinted Contigs (FPC) program has resulted in 13,040 contigs, consisting of 213,719 BACs, and 6,294 singleton BACs. The average contig size is 0.33 Mbp and the N50 contig size is 0.62 Mbp. WGP^TM^ technology has proved to provide a robust physical map of the pea genome, which would have been difficult to assemble using traditional restriction digestion based methods. This sequence-based physical map will be useful to assemble the genome sequence of pea. Additionally, the 1.1 million WGP tags will support efficient assignment of sequence scaffolds to the BAC clones, and thus an efficient sequencing of BAC pools with targeted genome regions of interest.

## Introduction

Field pea (*Pisum sativum* L.) is an important grain legume crop, which was domesticated ~7000 years ago (Abbo et al., 2010; Ambrose et al., 1995). The crop is valuable both for human nutrition and as animal feed. Gregor Mendel, the father of genetics, used pea as a model plant to uncover the fundamental principles of inheritance mainly because of the easily observable phenotypes and genotypes. However, understanding of quantitative traits and use of genomic tools for breeding is partly restricted by the large expected genome size of 3,947 to 4,397 Mbp/1C (Arumuganathan & Earle, 1991) and the occurrence of highly repetitive sequences in the pea genome. It is estimated that ~85% of the pea genome is of repetitive sequences (Murray et al., 1978). The majority of pea repetitive DNA is made of LTR-retrotransposons, which alone were estimated to contribute to 20-33% of the genome (Macas et al., 2007). In the current study, we have undertaken construction of a sequence-based physical map of pea to address the challenge in the assembly of these repetitive sequences and overcome the shortcomings of traditional restriction digestion based physical maps.

Whole genome profiling (WGP) is a sequence-based physical mapping technology for construction of bacterial artificial chromosome (BAC) contigs of complex genomes (van Oevern et al., 2011). WGP technology is based on generation of short sequence tags from terminal ends of restriction fragments of individual BAC clones, followed by assembly of BAC clones into contigs based on shared regions containing identical sequence tags. WGP is designed based on the use of sequence tags generated by next generation sequencing (NGS) and is a powerful alternative to traditional DNA fingerprinting based physical mapping technologies, and also simultaneously generates a partial genome sequence. Two-dimensional or multi-dimensional BAC clone pooling is an effective strategy for DNA preparation and sequencing to reduce the costs of sample preparation. The sequence-based physical map also provides information for localization of genes that are difficult to find through positional cloning. WGP was initially tested in *Arabidopsis thaliana* by using ~6,100 BAC clones and the assembly order of BAC contigs was verified with the genome sequence, wherein 98% of the BAC clones were assembled correctly (van Oevern et al., 2011). Following this validation, WGP was used to generate sequence-based physical maps and genome assembly of ~30 crop species (Ariyadasa & Stein, 2012; Sierro et al., 2013). WGP has been used for generation of physical maps of some individual wheat chromosomes, whose sequences are highly complex and repetitive (Philippe et al., 2012; Poursarebani et al., 2014). Recently, WGP technology was adopted by the International Wheat Genome Sequencing Consortium to generate new sequence information that will improve the quality and utility of physical maps for 15 chromosomes (www.wheatgenome.org). To address the challenges of accurate assembly of the massive and complex pea genome, we as part of international pea genome sequencing consortium adopted in the current study the WGP technology for assembly of pea BAC clones into a physical map.

## Materials and Methods

### BAC libraries

A total of 295,680 BAC clones derived from pea cv. Cameor available at the CNRGS, Toulouse, France, with an average insert size of 95 Kb and approximately 6.7-fold genome coverage were used to construct a sequence-based physical map (http://cnrgv.toulouse.inra.fr/layout/set/print/Library/Pea).

### Whole genome profiling (WGP)

#### Generation of BAC sequence tags

The BAC clones were subjected to WGP as described by van Oeveren et al. (2011). Pooling of BAC clones and DNA extraction was done by Amplicon Express (Pullman, WA). BAC clones stored in 384-well plates were pooled in a three-dimensional format, into row, column, and split-box pools, with each pool type consisting of 48, 48 and 64 clones, respectively. Illumina grade BAC DNA (high concentration and low *E.coli*) was extracted from the pooled BAC clones using an optimized alkaline lysis method. The DNA was digested with *Hind*III and *Mse*I restriction enzymes, ligated with Illumina adaptor sequences containing barcode sequences as sample identification tags and were PCR amplified. The PCR products were pooled, cluster amplified and amplicons were then sequenced from the *Hind*III restriction site end using the Illumina HiSeq2000 with 100 nt read length. The reads were processed for identification of barcodes and assigned to BAC pools followed by deconvolution, a process to assign sequence reads as WGP tags to individual BAC clones. Deconvolution was successful when the WGP tag was detected in exactly one of each of the three dimensions of the BAC pools. WGP tags were filtered for sequencing quality and used for contig analysis.

#### Physical map construction

A total of 825 million sequence tags were generated by WGP, of which 1.11 million tags were unique (Table S1) and corresponded to 220,013 BACs (Table S2). The unique sequence tags were used for construction of the physical map. These sequence tagged BACs were used to generate SuperBACs, by grouping all individual BACs with 75% or more similarity, using an improved version of Fingerprinted Contigs Software (FPC; Keygene^TM^). FPC was initially developed for analyzing BAC restriction fragment based fingerprint data (Soderlund et al., 1997), and the improved version is capable of processing sequence-based BAC fingerprint data. WGP tags from all the grouped BACs were assigned to the SuperBACs. WGP tags were converted into numbers to yield pseudo restriction fragment sizes for analysis using FPC to generate contigs based on BAC clone overlap. The genome coverage of BAC clones, mean contig size, and N_50_ contig size were calculated in million base pairs (Mbp) by multiplying FPC band units and the mean distance between two WGP tags.

## Results

### WGP tag generation

Multi-dimensional pooling of the 295,680 BAC clones and sequencing the ends of restriction fragments of pooled DNA generated 825 million deconvolutable reads, which constituted 45.5% of the total number of 1814 million high quality reads sequenced (Table 1). The deconvolutable reads yielded 1.11 million unique WGP tags and the average number of reads per tag was 96.6. The first 51 nucleotide sequence of the unique sequence tags are presented in Table S1. These WGP tags were tagged to 220,013 BACs (Table S2) with an average of 28.7 tags generated per BAC.

**Table 1.** Summary of whole genome profiling (WGP) input parameters and sequence data processing.

| WGP parameter |  |
| --- | --- |
| No. of BACs tested | 295,680 |
| Genome equivalents BACs tested | 6.7 |
| Enzyme combination WGP fragments | *HindIII/MseI* |
| No. of high-quality reads produced | 1814.8 million |
| No. of deconvolvable reads | 825.2 million |
| Percent deconvolvable reads | 45.5 |
| No. of unique WGP tags (FPC ready) | 1,108,689 |
| No. of tagged BACs (FPC ready) | 220,013 |
| Percent tagged BACs (FPC ready) | 84.6% |
| Average No. of WGP tags/BAC | 28.7 |
| Average No. of reads/tag | 96.6 |

### Physical map construction

The WGP tag data of 1.11 million tags tagged to 220,013 BAC clones was used to assemble individual BAC clones into contigs and superBACs using the modified FPC software (Keygene N.V.), capable of processing sequence-based BAC fingerprint data instead of fragment mobility information as used in the original FPC software (Soderlund et al., 1997). A cut-off value of E^−50^ was used initially to assemble the contigs. The cut-off value was reduced step-by-step and a final cut-off value of E^−01^ has resulted in 13,040 BAC contigs and 6294 BAC singletons. The number of BACs in each of the 13,040 contigs was listed in Table S3 and the BACs in each contig were listed in Table S4. The estimated N_50_ contig size was 42 BACs and average contig size was 0.329 Mbp. As an example, Fig. 1. shows the largest contig in the assembly, selected based on number of BACs and tags. The BACs are ordered to their positon in the contig. Horizontal lines indicate relative BAC length and positioning of the lines indicates relative position and degree of overlap between BACs. In Fig. 1 (A) Only non-buried BACs are shown, i.e., BACs which overlap with another BAC in the contig are not displayed, while Fig. 1 (B) shows part of the same contig but with all the buried BACs included.

**FIGURE 1.**
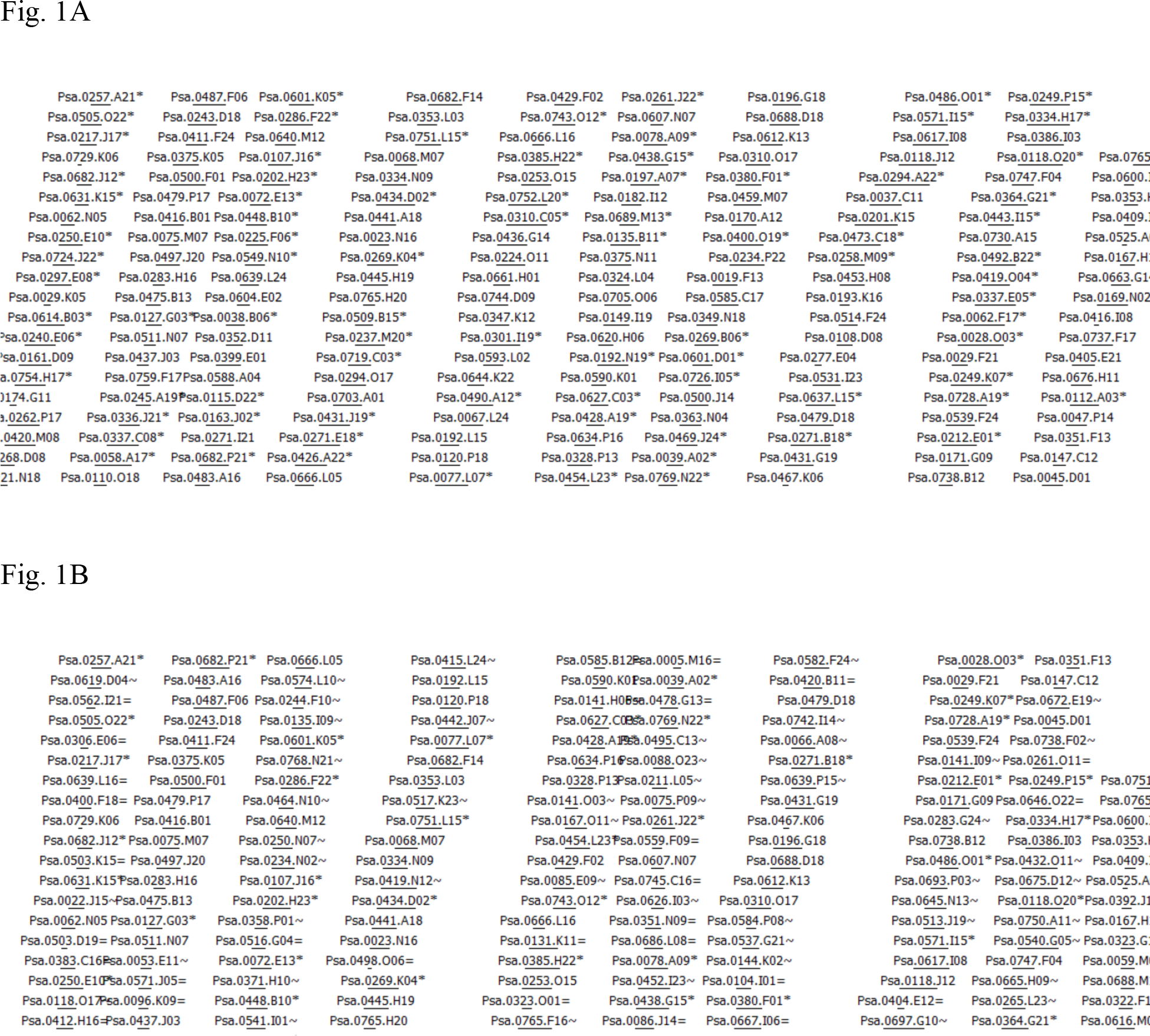
The largest contig in the assembly based on number of BACs and tags. The BACs are ordered to their positon in the contig. Horizontal lines indicate relative BAC length and positioning of the lines indicates relative position and degree of overlap between BACs. **(A)** Only non-buried BACs are shown, i.e., a semi-minimal tiling path, meaning that BACs which overlap largely with another BAC in the contig are not displayed. BACs indicated with a * indicate the presence of one or more buried BACs at this position. **(B)** Part of the same contig in the assembly showing all the buried BACs. Buried BACs are marked with = or ~, where = means identical and ~ means nearly identical. Note: The figures are in Consensus Band (CB) units; the length of the entire contig is 1532 CB units.

The estimated genome coverage was 4294 Mbp, which is 100% of the total estimated size of the pea genome (Table 2). After the deconvolution and filtering of the WGP tags, 27.7% of the BAC clones sequenced were not represented in the contig assembly. The parameters of physical map assembly are presented in Table 2.

**Table 2.** Whole genome profiling (WGP) metrics for the pea physical map construction using a 50 nt tag length and standard stringency.

| WGP parameter | Standard stringency WGP map assembly (E50<sup>-01</sup>) |
| --- | --- |
| No. of BACs in FPC | 220,013 |
| No. of contigs | 13,040 |
| No. of BACs in contigs | 213,719 |
| Percent of BACs FPC ready | 97% |
| No. of singleton BACs | 6,294 |
| Average contig size (BACs)<sup>1</sup> | 16.4 |
| N50 contig size (BACs)<sup>2</sup> | 42 |
| Average contig size (Mbp)<sup>3</sup> | 0.33 |
| N50 contig size (Mbp)<sup>4</sup> | 0.62 |
| Mbp coverage (%)<sup>5</sup> | 4,294 (100%) |

- 1) This is the mean number of BACs per contig. - 2) This number indicates that more than 50% of the contig coverage comprises contigs with at least this number of BACs - 3) This number is the mean contig size in million base pairs - 4) This number indicates that more than 50% of the contig coverage comprises at least this number of million base pairs - 5) The coverage estimate is based upon multiplication of FPC band units of all contigs with the estimated average distance between two tags. Due to this multiplication, the accuracy of the estimated average distance between two tags has a large impact on the result.

## Discussion

The two major steps involved in traditional physical map construction, restriction digestion-based fingerprinting several-fold genome equivalents of BAC clones, and their assembly into contigs, are highly intensive and error prone for a genome as large as pea. Several improvements have been made in BAC fingerprinting techniques (Luo et al., 2003) and contig assembly (Frenkel et al., 2010). The introduction of sequence-based WGP technology for physical map construction has made it possible to tag a large number of BAC clones based on short reads generated on NGS platforms and increase the accuracy of contig assembly (van Oeveren et al., 2011). This technology is particularly useful for large genomes with an abundance of repetitive DNA. Comparison of WGP sequence tags may also provide important biological information such as determination of ancestral origin of polyploids (Sierro et al., 2013).

The size of the current WGP-based physical map assembly corresponded with the estimated genome size of pea. The significance of this research includes the use of a large number of BAC clones, ~220,000, in WGP assembly and building a contig assembly near the estimated genome size of 4.2 GB, considering the high proportion of repetitive sequences. The parameters of the pea physical map assembly developed here are comparable to WGP-based physical maps of other crops, i.e., the average number of WGP tags per BAC clone (28.7) generated in this study and the percent of BAC clones represented in the contig assembly (72.3%) were comparable with WGP profiling of other complex genomes such as wheat (Poursarebani et al., 2014). Three contigs per Mbp were detected in the current physical map, in comparison to 2.2, 2.6 and 3.1 contigs per Mbp reported in tobacco (Sierro et al., 2013), tomato, and potato (De Boer et al., 2011), respectively. In the pea physical map assembly, the average number of BACs per contig is 16.4 and the average contig size is 0.33 Mbp in comparison to 34 BACs and 0.46 Mbp in tobacco (Sierro et al., 2013).

In this research, we have constructed a high quality physical map of pea based on WGP with the assembly parameters comparable to WGP assembly of other crops. Since the map is based on sequenced DNA tags, the physical map provides the skeleton framework for anchoring the genome sequence to obtain a high quality reference genome sequence to explore the genes governing traits and to study the genome features. The recent improvements of optical mapping of genomes in nanochannel arrays (Bionano) (Lam et al., 2012) and `Chicago` method based on *in vitro* reconstituted chromatin (Putnam et al., 2016) are further advancements to support physical mapping and sequence assembly in complex genomes and provide substantial improvement in the N_50_ contig size. Using the Bionano approach, Stankova et al. (2016) obtained contigs of the short arm of chromosome 7D (7DS; 381Mb) of bread wheat, with a N_50_ value of 1.3Mb, and identified ~800kb array of tandem repeats.

We have provided information of all the WGP tags in Supplementary Table S1 and the BACs corresponding to these tags are shown in Supplementary Table S2. The map is accessible through the .FPC file (Supplementary file F1), and users can view it in FPC output format, by using FPC software. This information will assist users to navigate and identify the BAC clones of their interest. The international consortium for pea genome sequencing is using the WGP-based physical map in conjunction with Bionano optical mapping to anchor and improve the complex genome sequence of pea.

## Conflict of Interest Disclosure

The authors declare that there are no conflicts of interest related to the manuscript.

## Acknowledgements

The study was funded by Saskatchewan Pulse Growers (SPG).

## Author Contributions

TW, BT, JB and EV designed the study. JV and KL performed the sequence and FPC analysis. HB and AB provided the BACs. KKG drafted the manuscript. RVBL contributed to data analysis. All authors contributed to the manuscript review.

---

**Supplemental Table S1** Unique sequence tags identified by sequencing ends of restriction fragments of 295,680 BAC clones.

**Supplemental Table S2** BAC clones corresponding to the unique sequence tags identified by sequencing ends of restriction fragments of 295,680 BAC clones.

**Supplemental Table S3** Number of BAC clones in each contig built based on the sequence similarities of unique sequence tags.

**Supplemental Table S4** Distribution of BAC clones in contigs built based on the sequence similarities of unique sequence tags.

## References

Abbo, S., Lev-Yadun, S., and Gopher, A. (2010). Agricultural origins: Centers and noncenters; a near eastern reappraisal. Crit. Rev. Plant Sci. 29(5), 317–328.

Ambrose, M. J. (1995). From Near East centre of origin the prized pea migrates throughout world. Diversity 11(1–2), 118–119.

Ariyadasa, R., and Stein, N. (2012). Advances in BAC-based physical mapping and map integration strategies in plants. J. Biomed. Biotechnol. doi:10.1155/2012/184854.

Arumuganathan, K., and Earle, E. D. (1991). Nuclear DNA content of some important plant species. Plant Mol. Biol. Report. 9(3), 208–219.

De Boer, J. M., Borm, T. J. A., Jesse, T., Brugmans, B., Tang, X., and Bryan G. J., et al. (2011). A hybrid BAC physical map of potato: a framework for sequencing a heterozygous genome. BMC Genomics. 12, 594.

Frenkel, Z., Paux, E., Mester, D., Feuillet, C., and Korol, A. (2010). LTC: A novel algorithm to improve the efficiency of contig assembly for physical mapping in complex genomes. BMC Bioinformatics. 11, 584.

Lam, E. T., Hastie, A., Lin, C., Ehrlich, D., Das, S. K., Austin, M. D., et al. (2012). Genome mapping on nanochannel arrays for structural variation analysis and sequence assembly. Nat. Biotechnol. 30(8), 771–776.

Luo, M. C., Thomas, C., You, F. M., Hsiao, J., Ouyang, S., Buell, C. R., et al. (2003). High-throughput fingerprinting of bacterial artificial chromosomes using the snapshot labeling kit and sizing of restriction fragments by capillary electrophoresis. Genomics. 82(3), 378–389.

Macas J., Neumann, P., and Navratilova, A. (2007). Repetitive DNA in the pea (Pisum sativum L.) genome: comprehensive characterization using 454 sequencing and comparison to soybean and Medicago truncatula. BMC Genomics. 8, 427.

Murray, M. G., Cuellar, R. E., and Thompson, W. F. (1978). DNA sequence organization in the pea genome. Biochemistry. 17(26), 5781–5790.

Philippe, R., Choulet, F., Paux, E., Oeveren, J. V., Tang, J., Wittenberg, A. H. J., et al. (2012). Whole Genome Profiling provides a robust framework for physical mapping and sequencing in the highly complex and repetitive wheat genome. BMC Genomics. 13, 47.

Poursarebani, N., Nussbaumer, T., Simkova, H., Safar, J., Witsenboer, H., van Oeveren, J., et al. (2014). Whole-genome profiling and shotgun sequencing delivers an anchored, gene-decorated, physical map assembly of bread wheat chromosome 6A. Plant J. 79(2), 334–347.

Putnam, N. H., O’Connell, B. L., Stites, J. C., Rice, B. J., Blanchette, M., Calef, R., et al. (2016). Chromosome-scale shotgun assembly using an in vitro method for long-range linkage. Genome Res. 26(3), 342–350.

Sierro, N., van Oeveren, J., van Eijk, M. J. T., Martin, F., Stormo, K. E., Peitsch, M. C., Ivanov, N. V. (2013). Whole genome profiling physical map and ancestral annotation of tobacco Hicks Broadleaf. Plant J. 75(5), 880–889.

Soderlund, C., Longden, I., and Mott, R. (1997). FPC: a system for building contigs from restriction fingerprinted clones. Comput. Appl. Biosci. 13(5), 523–535.

Staňková, H., Hastie, A. R., Chan, S., Vrana, J., Tulpova, Z., Kubalakova, M., et al. (2016). BioNano genome mapping of individual chromosomes supports physical mapping and sequence assembly in complex plant genomes. Plant Biotech J. 14(7), 1523–1531.

van Oeveren, J., de Ruiter, M., Jesse, T., van der Poel, H., Tang, J., Yalcin, F., et al. (2011). Sequence-based physical mapping of complex genomes by whole genome profiling. Genome Res. 21(4), 618–625.

